# Maternal allergic sensitization affects the T cell modulatory capacity and molecular cargo of milk derived extracellular vesicles

**DOI:** 10.1101/2023.08.11.553032

**Authors:** Martijn J.C. van Herwijnen, Alberta Giovanazzi, Marijke I. Zonneveld, Joaquín J. Maqueda, Marije Kleinjan, Soenita Goerdayal, Franziska Völlmy, Arianne van Bruggen - de Haan, Tom A.P. Driedonks, Ger J.A. Arkesteijn, Ruurd M. van Elburg, Gerbrich N. van der Meulen, Johan Garssen, Carla Oliveira, A.F. Maarten Altelaar, Peter A.C. ’t Hoen, Frank A. Redegeld, Esther N.M. Nolte – ’t Hoen, Marca H.M. Wauben

## Abstract

**Background:** Human milk extracellular vesicles (EVs) affect various cell types in the gastrointestinal tract, including T cells, and play a role in the development of the newborn’s immune system by delivering specific molecular cargo to target cells. Although maternal allergic sensitization alters the composition of milk, it is unknown whether this impacts the function of milk EVs. Therefore, we analyzed the T cell modulatory capacity and compared the protein and miRNA cargoes of EVs from milk of allergic and non-allergic mothers.

**Methods:** EVs were isolated from human milk from allergic and non-allergic donors by differential centrifugation, density gradient floatation and size exclusion chromatography. Functional modulation of primary human CD4+ T cells by EVs was assessed *in vitro*. Proteomic analysis and small RNA sequencing was performed on milk EVs to evaluate protein and miRNA abundance and to identify cellular targets of this EV cargo in relevant T cell signaling pathways.

**Results:** T cell proliferation, activation and cytokine production were suppressed in the presence of milk EVs. Remarkably, milk EVs from allergic mothers inhibited T cell activation to a lesser extent than EVs from non-allergic mothers. Integrative multi-omics analysis identified EV cargo of which the cellular targets could be linked to differential modulation of T cell activation-associated processes .

**Conclusions:** Milk EVs from non-allergic mothers are stronger inhibitors of T cell activation compared to milk EVs from allergic mothers. This altered functionality might be linked to changes in miRNA and protein cargo that modulate T cell signaling pathways in an integrative manner.

## Introduction

Allergic disease is an escalating health hazard and contributes substantially to child morbidity [Prescott]. Despite extensive experimental and epidemiological research, the impact of maternal sensitization on allergy development in breastfed infants remains unclear [Hamada, Leme, Matson, Lodge, Odijk, Verhasselt]. Although maternal allergy has been linked to an altered milk composition [Prokesová, Laiho, Kuitunen, Hettinga, Böttcher, Torregrosa Paredes], little is known about the functional consequences due to this different composition. Therefore it is crucial to study the immune modulatory mechanisms of different milk components that contribute to the development of the immune system. Cell-derived extracellular vesicles (EVs) have been identified as bio-active components present in human milk [Admyre, Zonneveld 2021]. EVs are nano-sized lipid bilayer enclosed structures involved in intercellular communication and playing pivotal roles in immune regulation [Buzas, Théry]. Milk EVs harbor proteins involved in development and immune regulation [van Herwijnen 2016] and contain miRNAs, which are potent regulators of cell signaling [van Herwijnen 2018]. As human milk EVs have a strong CD4+ T cell modulatory potential [Admyre, Zonneveld 2021] and CD4+ T cells are instrumental in the development of allergic disease, we investigated whether milk EVs from allergic and non-allergic mothers differentially affect CD4+ T cell responses. We subsequently performed proteomics and small RNA (sRNA) sequencing analysis of purified milk EVs to evaluate changes in EV cargoes related to the maternal allergic status and used an integrative omics approach to identify cellular targets that can modulate T cell signaling pathways.

## Materials & Methods

### Donor selection and classification of allergic and non-allergic subjects

This study was performed as part of the Comparison of Human Milk Extracellular Vesicles in Allergic and Non-allergic Mothers (ACCESS) study (NL 47426.099.14; RTPO 914). Donors were lactating women of good general health, aged 18 years or above, who had given birth to a full-term newborn via vaginal delivery. Exclusion criteria were: use of immune suppressive or immune modulatory medication, immune-related diseases (other than allergy), severe atopic dermatitis, smoking, drug abuse, pre-eclampsia, and alcohol consumption during lactation. Milk samples were provided during routine hospital check-ups. Donors were classified as ‘allergic’ if total serum IgE ≥ 50 kU/L and/or specific IgE was detected for grass pollen, tree pollen, house dust mite, cat dander, or dog dander by positive Phadiatop assay (≥ 0.35 kU/L; Thermo Scientific, Uppsala, Sweden). Non-allergic donors had no history and symptoms of allergy (absence of any atopy related condition, including diagnosed allergic asthma and diagnosed atopic dermatitis), total serum IgE < 50 kU/L and Phadiatop assay was negative (< 0.35 kU/L). The summary with donor information is provided in Table 1. Individual donor information is provided in Supplementary Table 1. All donors signed an informed consent form and the study was approved by the local medical ethics committee.

**Table 1:** Summary of the baseline characteristics of the participating donors. Table lists various donor information and whether donors were classified as non-allergic or allergic. Significance was calculated using unpaired two-tailed t test for normally distributed data, Mann-Whitney test for nonparametric data, and Fisher exact for dichotomous data.

| **Baseline characteristics** | **non-allergic**<br>**(n=30)** | **allergic**<br>**(n=31)** | **P value** |
| --- | --- | --- | --- |
| maternal age (y, mean ± SD) | 31.5 ± 3.7 | 31.6 ± 3.5 | 0,876 |
| lactational stage (wk, mean ± SD) | 7.0 ± 2.3 | 6.4 ± 1.9 | 0,313 |
| infant sex (male/female) | 14/16 | 17/14 | 0,612 |
| parity (mean number of infants, mean ± SD) | 1.80 ± 1.1 | 1.74 ± 0.9 | 0,824 |
| **maternal history of inhalation allergy (n, %)** | **3 (10.0%)** | **20 (64.5%)** | **<0,0001** |
| **maternal total IgE (mean, kU/L ± SD)** | **18.9 ± 14.8** | **282.5 ± 881.9** | **<0,0001** |

### Follow up study on the health status of the infants

Participating mothers received questionnaires 1 year after enrolment in the study. The questionnaire was designed to obtain data about the health status of the infants during the 1^st^ year of life and contained the following sections: general health (including questions about perception of health), infectious diseases (including questions about sneezing or a stuffy nose) and allergic diseases (including questions about itching rash and its frequency and its location, losing sleep over itching rash; occurrence of wheezing, its frequency, losing sleep over wheezing; dry coughs without having a cold; low-pitched wheezes or coarse crackles; shortness of breath; diagnosis of asthma; itching eyes) (Supplementary Table 2).

### Milk collection and isolation of milk EVs and EV-depleted control for functional T cell assays

Collection of human milk and isolation of EVs and EV-depleted procedural controls were done as previously described [Zonneveld 2021] using differential centrifugation, density gradient separation and size exclusion chromatography. Importantly, the concentration of EVs used in T cell assays resembles the physiological concentration of EVs in milk as 7 ml of purified EV sample was obtained from 6.5 ml milk supernatant. We previously validated this isolation protocol by Western blot analysis and nanoparticle tracking analysis NTA [Zonneveld 2021]. Data are available in the EV-TRACK knowledgebase (EV-TRACK ID: EV200007) [Van Deun].

### CD4+ T cell isolation and stimulation

T cells were isolated from peripheral blood mononuclear cells (PBMC) as previously described [Zonneveld 2021]. Purified CD4+ T cells were seeded at 0.5*10^6 cells/ml in plates coated with 0.5 μg/ml anti-CD3 (CLB-T3/4.E, 1XE) and 70 ng/ml soluble anti-CD28 (CLB-CD28/1, 15E8; both from Sanquin). Cells were cultured in a total volume of 1 ml consisting of medium only, or of 750 μl EVs or EV-depleted control + 250 µl medium for the indicated amount of time.

### Multiplex cytokine analysis

Supernatants from stimulated T cells were harvested on day 2 and day 6 depending on peak of cytokine production and analyzed using the LEGENDplex^TM^ human Thelper 1/2/9/17 multiplex kit (BioLegend). Beads were acquired on a BD Canto II (BD Bioscience) and analyzed with LEGENDplex^TM^ V7.0.

### Flow cytometry

Proliferation of stimulated CD4+ T cells was assessed by labeling cells with 2 μM CellTrace Violet (Invitrogen) prior to culture. Following culture, cells were harvested and stained with fluorescent conjugated antibodies. Antibodies used in this study: CD25-Alexa488 (eBioscience), CD3-PE-Cy7 and CD4-PerCP-Cy5.5 (all from BioLegend). Cells were acquired on a BD Canto II (BD Bioscience), and analyzed by FlowJo (V10.1).

### RT^2^ Profiler PCR array

Real-time quantitative PCR (RT-qPCR) was done as previously described [Zonneveld 2021] using the Human T cell Tolerance & Anergy RT^2^ Profiler PCR array (Qiagen). The relative expression levels of each gene were normalized using 4 reference genes (B2M, GAPDH, HPRT1, RPLP0).

### EV isolation for Liquid chromatography tandem mass spectrometry (LC-MS/MS), sRNA sequencing and high resolution flow cytometry

Milk EVs were isolated from 6.5 ml (for proteomics and high resolution flow cytometry) or 3.5-4 ml (for sRNA sequencing) 10,000 g milk supernatant as previously described [van Herwijnen 2016], using differential centrifugation and sucrose density gradient separation. Data on EV characterization are available in the EV-TRACK knowledgebase (EV-TRACK ID: EV160000) [Van Deun]. EV-enriched fractions from sucrose gradient isolation were diluted with PBS (Invitrogen) and pelleted at 192,000 g using a SW40Ti rotor. EV pellets were either frozen directly at −80°C until high resolution flow cytometry and protein extraction for LC-MS/MS, or resuspended in 700 µl Qiazol and frozen at −80°C until sRNA isolation for sequencing.

### EV protein extraction, digestion and high resolution LC-MS/MS mass spectrometry

EV protein extraction was performed as previously described [Van Herwijnen 2016]. Protein digestion was performed with filter aided sample preparation (FASP) [Wiśniewski], with a buffer containing 8 M urea in 1 mM Tris-HCl at pH 8.0 and pH 8.5; using filters with cutoff of 10 kD (Merck Millipore). Proteins were reduced with 10 mM DTT alkylated with 200 mM iodo acetamide (both from Sigma-Aldrich) and digested for 4 hrs with 2 mg/ml LysC (Wako Chemicals GmbH) in a ratio of 1:50 and O/N with 0.1 µg/µl trypsin (Promega) in a ratio of 1:50. The digested samples were desalted using Oasis HLB 96-well µelution plate (Waters Chromatography B.V.). Finally, protein samples were dried and reconstituted in 40 µl of 10% formic acid/5% DMSO (Sigma-Aldrich). Samples were analyzed as previously described [Van Herwijnen 2016] with a QExactive Plus instrument (Thermo Scientific) connected to an Agilent 1260 Infinity LC system equipped with a 20 mm x 100 µm ID Reprosil C18 (Dr Maisch HPLC GmbH) trap column and a 450 mm x 75 µm ID Poroshell C18 analytical column (Agilent Technologies). Small deviations from Van Herwijnen et al. [Van Herwijnen 2016] were a flow rate of 5 µl/min for 10 min during trapping and elution of peptides using a passively split flow of 180 nl/min (120 min LC method). For the QExactive analysis the 20 most intense ions in the survey scan (375 to 1600 m/z, resolution 35,000, AGC target 3e6) were fragmented in the ion trap and were subjected to HCD fragmentation (resolution 17,500, AGC target 5e4), with the normalized collision energy set to 25% for HCD. The signal threshold for triggering an MS/MS event was set to 500 counts. Dynamic exclusion was enabled (exclusion size list 500, exclusion duration 18 s). The mass spectrometry data will be deposited to the ProteomeXchange Consortium via the PRIDE [Vizcaino] repository.

### LC-MS/MS data processing

MS raw data were processed with MaxQuant (version 1.6.17.0). Generated peak lists were searched against Swissprot *Homo sapiens* database, (April 2021, 20,407 entries) supplemented with frequently observed contaminants using Andromeda, MaxQuant’s built-in search engine. Trypsin was chosen with two missed cleavages allowed. Oxidation (M) and acetylation (protein N-term) were set as variable modifications and carbamidomethylation (C) was set as a fixed modification. The searches were otherwise performed using default parameters, resulting in 1% false discovery rate (FDR).

### Isolation of EV-sRNA, library preparation and sRNA sequencing

RNA was isolated using the miRNeasy micro kit (Qiagen) according to manufacturer’s instructions. Quality and quantity of the RNA samples was assessed using Agilent 2100 Bioanalyzer pico-RNA chips (Agilent Technologies Netherlands B.V.).

Input quantity of sRNA for library preparation was based on equal volume of purified milk EVs and ranged between 10-150 ng of sRNA per sample. The Illumina® Truseq small RNA Sample Prep Kit was used to process the samples according to the kit-specific guidelines. Briefly, adapters were ligated to each end of sRNA, sRNA was converted into cDNA, and PCR amplification was performed. The quality and yield after sample preparation was measured with the Fragment Analyzer. Small RNA sequencing was done on the Illumina NextSeq 500 platform with 75 bp single-end sequencing. Samples were sequenced in 1 flow cell with 4 lanes. Samples with less than 7 million reads were sequenced in a second flow cell and data from 2 flow cells were combined.

### Small RNA sequencing data processing

For each sample, 4 lanes were combined in a single fastq file. Samples sequenced in 2 flow cells were combined. In total 20 fastq files from non-allergic donors and 20 from allergic donors were obtained. FastQC (version 0.11.5) [Andrews] was used to perform quality control checks on sequencing data. Remaining TruSeq small RNA adapter sequences were checked and clipped from all reads at the 3’ end using cutadapt (version 2.8, with settings of “-a TGGAATTCTCGGGTGCCAAGGAACTCCAGTCAC --error-rate 0.1 --times 1 -m 15”) [Martin]. Remaining adapters were removed from the 5’ end of the reads using cutadapt (version 2.8, with settings of “-g CGACAGGTTCAGAGTTCTACAGTCCGACGATC --error-rate 0.1 -- times 1 -m 15) [Martin]. Low quality bases were trimmed from all sequence reads using sickle (version 1.33, with settings of “-t sanger -l 15”) [Joshi]. All sequence reads shorter than 15 bases were discarded. The processed reads were aligned to human reference genome GRCh38 from Ensembl [Cunningham] (*Homo sapiens* primary assembly, soft-masked) and the number of aligned reads per annotated sRNA was calculated by Manatee algorithm (version 1.2) [Handzlik] applying bowtie to align reads (version 1.0.1, with settings of “--best --strata -m 50 -k 3 -v 3”) [Langmead], allowing for alignments to a maximum of 50 loci and reporting 3 of them, with equal alignment scores, and a maximum of 3 mismatches. Manatee counts output was rounded and used as raw expression data. The small RNA annotation track was obtained from miRBase (version 22.1) [Kozomara], GtRNAdb (version 2.0, January 2016) [Chan] and Ensembl 99 [Cunningham] in order to analyse the miRNAs. Sequencing data were deposited in NCBI’s Gene Expression Omnibus (GEO) (GSE216498).

### Analysis of differential protein and miRNA expression

Proteins and miRNAs with at least 1 peptide spectral match (PSM) or 1 count per million (CPM) present in 60% of samples (3/5 for proteomics and 12/20 for sRNA analysis) from allergic or non-allergic were considered valid for analysis and used as input for differential expression analysis.

For proteins, the LFQ normalized intensities were used as quantitative input. Values were log2 transformed, and submitted to a two-sided Student’s t-test to assess significance (p-value < 0.05) between allergic and non-allergic groups. For miRNAs, significant differential expression between the libraries of allergic vs. non-allergic milk donors was tested using the edgeR package (version 3.32.0) [Robinson 2009] Data were normalized using the TMM method (weighted trimmed mean of M-values) [Robinson 2010]. A generalized linear model was fit after estimating the common, trended and tagwise dispersion. Due to adapter contamination, 2 batches were observed in our samples. Therefore, we included allergy status and batch as fixed effects in the model. The likelihood ratio test was used to evaluate differential sRNA expression between allergic and non-allergic donors. Differences in expression between conditions with |log2(fold-change)| > log2(1.5) and non-adjusted p-value < 0.05 were considered to be significant. In an additional analysis, proteins and miRNAs were selected when present in at least 60% in one condition and less than 60% in the other condition and significantly differed between allergic and non-allergic donors, based on the Mann-Whitney test using the non-normalized mean intensity of each protein and CPM of the miRNAs (p-value < 0.05).

### Target prediction and Gene Ontology (GO) analysis

In order to determine the cellular targets of the EV cargo in T cells, we analyzed the targets of the differentially expressed proteins and miRNAs. For differentially expressed proteins we used String (Version 11.5) [Szklarczyk], with active interaction sources set to ‘experiments’ only and confidence set to ‘default: medium confidence 0.400’ and analyzed no more than 50 interactors to identify potential cellular targets. For differentially expressed miRNAs, we used the miRNA-target interactions (MTI) miRTarBase (release 8.0) [Hsu] to select strong evidence targets only found by reporter assays (Functional MTI) to identify cellular targets. Next, only cellular targets from both proteins and miRNAs that have been experimentally identified in either naïve or CD28/CD3-activated T cells (DICE database, inclusion criterium: tpm≥1 in at least one category: naïve or activated) were used for GO analysis. The Cytoscape plugin ClueGO (version 2.5.7) [Shannon] was used to compute a list of significantly enriched GO terms (biological processes from all experimental evidences – EXP, IDA, IPI, IMP, IGI, IEP; GO tree interval 3-8, Go term included with min 3 genes as 4% total genes, κ score 0.7) linked to targets over-represented in non-allergic EVs and allergic EVs. A graphical representation of a network connecting these terms based on similarity in ontology was made, where nodes represent GO terms. The function of proteins associated to selected GO terms were searched for in the Uniprot database (release 2022_03) and their action was depicted in T cell pathway network models using Inkscape (version 1.2.1). Full target prediction and GO analysis can be found in Supplementary file 1.

### Statistical analysis

Data normality was assessed by the Shapiro–Wilk test and homogeneity of variances was confirmed with the use of the Brown–Forsythe test. Statistical analysis on baseline characteristics of participating donors and infant health was calculated using unpaired two-tailed t test for normally distributed data, Mann-Whitney test for nonparametric data, and Fisher exact for dichotomous data. Differences between the functional effects of EVs and EV-depleted samples and differences between allergic and non-allergic donors were analyzed simultaneously by repeated measures two-way ANOVA and Sidak’s multiple comparisons test. Differences of EV cargo between groups was determined by two-sided Student’s t-test and Mann-Whitney test. For GO-analysis on cellular targets Fisher Exact Test was used. Calculations were made in GraphPad Prism Software V8.3.0. Significance was defined as *p < 0.05, **p < 0.01, ***p < 0.001, and ****p < 0.0001.

## Results

### Differential modulation of CD4+ T cell activation by human milk EVs from non-allergic and allergic donors

In order to evaluate whether allergic sensitization impacts the T cell modulatory activity of EVs, activated CD4+ T cells were cultured in the presence of EVs and corresponding EV-depleted controls isolated from milk of allergic or non-allergic mothers. EVs significantly inhibited CD4+ T cell proliferation and CD25 expression, compared to their donor-matched EV-depleted controls (Figure 1). However, while milk EVs from non-allergic donors inhibited IL-6 and TNFα cytokine production, EVs derived from allergic donors did not reduce release of these pro-inflammatory cytokines. Moreover, there was a significant increase of CD4+ T cells expressing CD25 and producing IL-6 and TNFα upon exposure to EVs of allergic donors compared to EVs from non-allergic donors (Figure 1), indicating an sensitization-dependent modulation of T cell activation. Other cytokines tested showed no significant differences, as EVs from both groups inhibited overall cytokine secretion compared to EV-depleted controls (Supplementary Figure 1). Importantly, no differences were observed between EV-depleted controls of allergic or non-allergic mothers, indicating that only EVs are affected by allergic sensitization.

**Figure 1.**
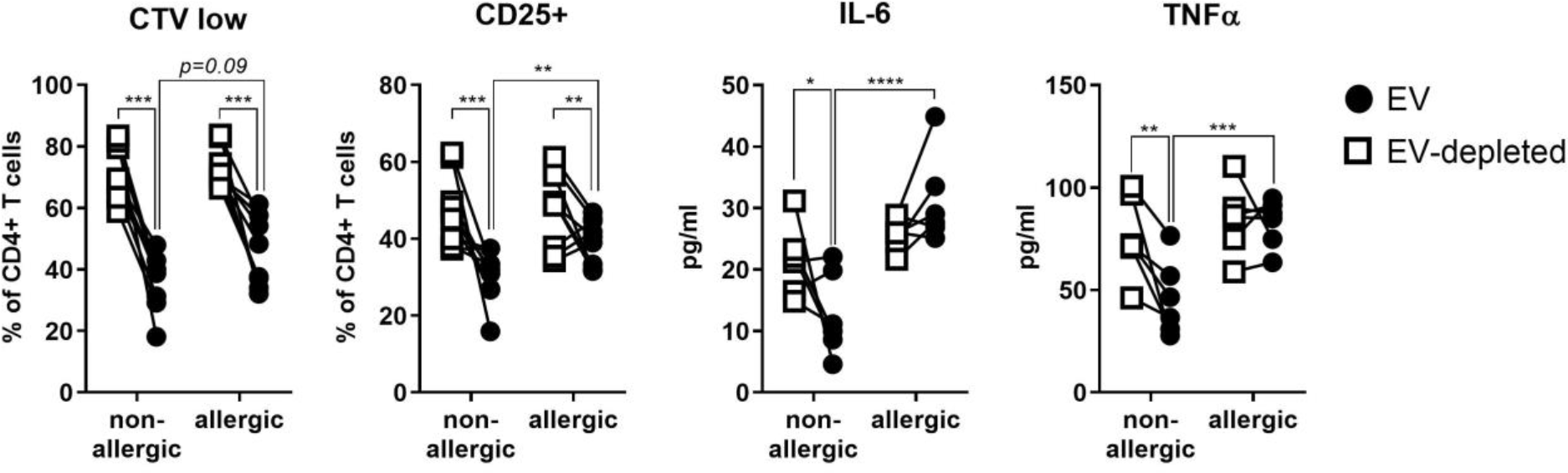
Inhibition of CD4+ T cell activation is reduced by milk EVs from allergic mothers. CD4+ T cells were stimulated with anti-CD3 and anti-CD28 in the presence of EVs or its EV-depleted control sample from allergic (n=3) or non-allergic (n=3) mothers. For flow cytometry, T cells from 3 different donors were labeled with CellTrace Violet (CTV), cultured for 6 days, harvested and stained with anti-CD25 and analyzed. Quantification of proliferation is shown for cells which had undergone one or more proliferation cycles (and were thus lower in CTV signal intensity than non-proliferating T cells). Expression of the activation marker CD25 is indicated as the percentage of CD4+ T cells having a positive staining for CD25. For cytokine production, T cells of 2 different donors were cultured for 48 hours after which supernatants were harvested and analyzed for IL-6 and TNFα. Graphs show the paired results obtained for each EV donor (●) and its EV-depleted control (□). Significance was calculated using repeated measures two-way ANOVA with Sidak’s multiple comparisons test. -values are defined as **p < 0.01, **p<0.01, and ****p < 0.0001.

Considering that maternal characteristics can influence milk composition, including EV numbers [Torregrosa Paredes] we found that the difference in T cell modulatory capacity could not be explained by a difference between donors as maternal age, lactational stage, distribution of infant sex and parity were similar (Table 1). However, when we determined EV numbers by high resolution flow cytometry, allergic mothers had less milk EVs compared to non-allergic mothers (supplementary Figure 2). When we normalized EV numbers in a functional comparison between a non-allergic donor to an allergic donor, the difference in T cell modulation remained, indicating that qualitative changes rather than quantitative changes in milk EVs cause a difference in T cell modulation (supplementary Figure 3). Therefore, we set out to investigate any changes in EV composition due to maternal sensitization.

### Comparison of protein and miRNA content of milk EVs from non-allergic and allergic mothers

We performed proteomics and sRNA sequencing on milk EVs of non-allergic or allergic mothers and identified 650 unique proteins and 1898 unique miRNAs in both groups of which 515 proteins and 714 miRNAs were present in at least 60% of the samples in either condition, which we used as a trusted set for differential expression analysis. For this, the average intensities of the proteins and CPM of the miRNAs were used as a measure of abundance, which were normalized to calculate the fold change (FC) in expression between allergic and non-allergic milk EVs. Proteins and miRNAs with a FC greater than 1.5 (log2FC >0.58) and smaller than 0.66 (log2FC <-0.58) and a non-adjusted *p*-value <0.05 (-log10*P* >1.3) were considered significantly differentially expressed.

We observed that the majority of proteins and miRNAs were not differentially expressed. However, we identified 3 enriched proteins in allergic donors (PGM1, SPP1 and TF) and 5 enriched proteins in non-allergic donors (C4BPA, CYB5, MUC15, RPS8 and RTN3), while for the miRNAs we identified 11 enriched miRNAs in allergic donors (miR-10a-3p, miR-10a-5p, miR-100-5p, miR-378g, miR-548ao-3p, miR-369-3p, miR-642a-5p, miR-940, miR-3173-5p, miR-4688, miR-6757-5p) and 15 enriched miRNAs in non-allergic donors (miR-142-3p, miR-142-5p, miR-223-3p, miR-338-3p, miR-556-5p, miR-582-5p, miR-889-3p, miR-1234-3p, miR-3614-5p, miR-3616-3p, miR-4459, miR-6726-3p, miR-6729-5p, miR-6853-3p, miR-7641) (Figure 2A: volcano plots, left panels). Albeit after multiple testing correction these individual proteins and miRNAs did not meet significance thresholds, we aimed to identify EV cargo as groups of proteins and/or miRNAs that are associated to the maternal allergic status.

**Figure 2.**
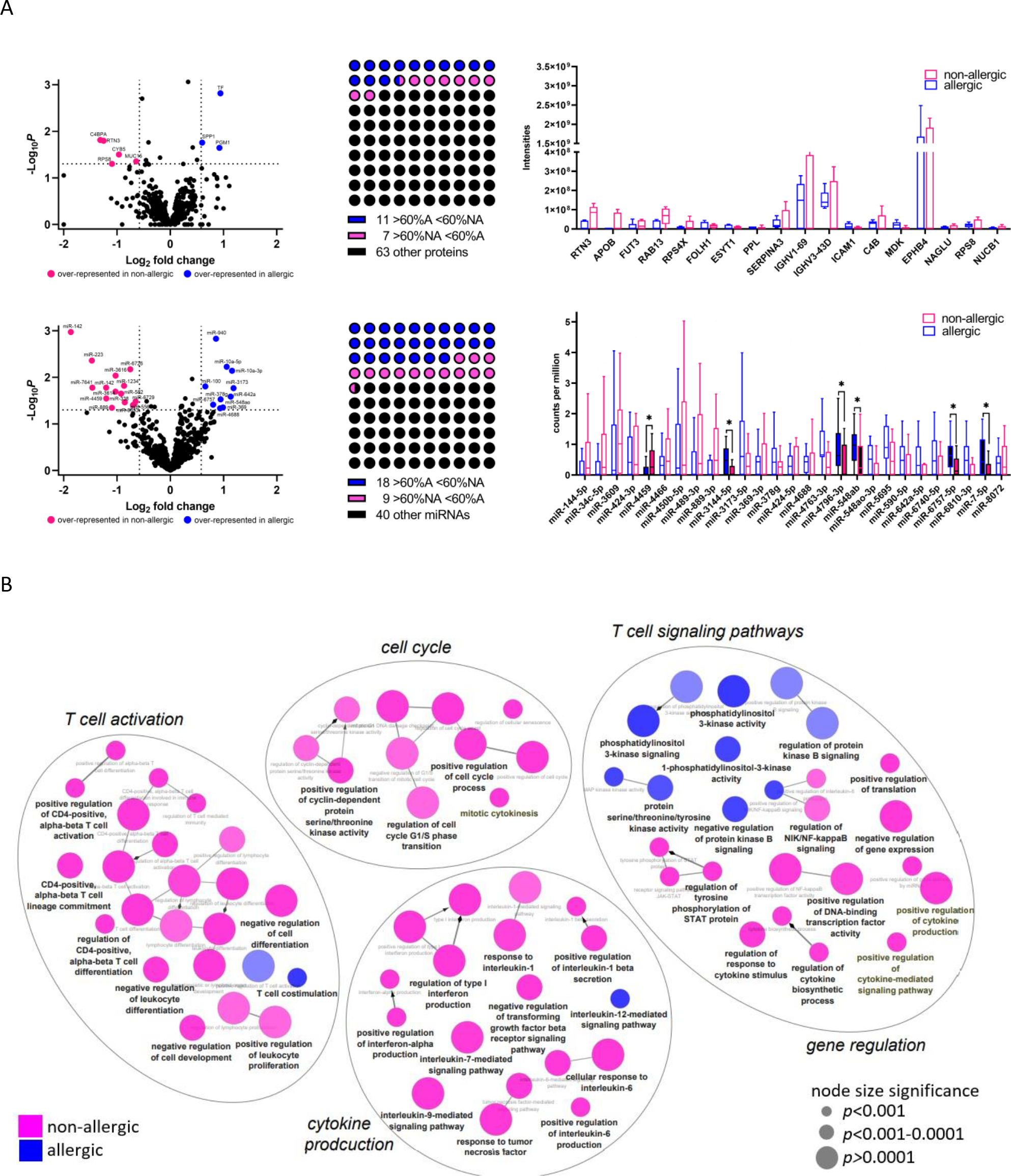
Omics approach to identify protein and miRNA content of EVs from allergic and non-allergic mothers. (A) Proteomics was performed on n=5 individual samples per group and sRNA sequencing on n=20 individual samples per group on purified EVs from milk of non-allergic or allergic mothers. Proteins are depicted in the upper part and miRNAs in the bottom part of figure A. We used the LFQ normalized intensities from the proteomics as quantitative input which were log2 transformed and submitted to a two-sided Student’s t-test to assess significance between allergic and non-allergic groups. For miRNAs, the mean log2 (CPM) – TMM normalized of each miRNA was calculated and likelihood ratio test was used to evaluate differential expression between allergic and non-allergic donors. A generalized linear model was fit after estimating the common, trended and tagwise dispersion (volcano plots, left panel). Differential expression, defined as differences in expression between conditions with |log2(fold-change)| > log2(1.5) and non-adjusted p-value < 0.05, is indicated as dotted line in the volcano plots. Additional differential expression analysis selected proteins and miRNAs in at least >60% samples from one condition and <60% in the other and a FC of at least 1.5 were selected (dotplots, middle panels). From these selected candidates, for each condition the mean CPM of each miRNA or mean intensity of each protein (no normalization applied, no dispersion corrected, no batch corrected) was calculated and Mann-Whitney test was applied to determine significant differences (box plots, right panel). (B) GO analysis of identified cellular targets depicted in a network. Color coding shows enrichment in GO terms of either non-allergic (pink) or allergic (blue). The size of the nodes reflects the statistical significance of the terms (as determined by Fisher Exact Test).

Therefore, we performed additional differential expression analysis as defined by FC >1.5 or FC <0.66 in at least 60% of the samples in one condition and less than 60% in the other condition with a p-value<0.05 without any normalization or corrections based on the raw data. Hereto, we first selected cargo with FC >1.5 or FC <0.66, which resulted in 81 proteins and 67 miRNAs (Figure 2A: dotplots, middle panels). Next, these proteins and miRNAs were analyzed for their presence in at least 60% in one condition and less than 60% in the other condition. This resulted in 11 proteins and 18 miRNAs more represented in the non-allergic group, and 7 proteins and 9 miRNAs more represented in the allergic group (Figure 2A: dotplots, middle panels). Finally these proteins and miRNAs were tested for significance (Mann-Whitney test) (Figure 2A: box plots, right panels). Although none of the proteins showed significant differences, 6 miRNAs were differentially expressed (without multiple testing correction) of which 5 were enriched in EVs derived from allergic donors (miR-7-5p, miR-548-ab, miR-3144-5p, miR-4796-3p, miR-6757-5p) and 1 in EVs from non-allergic donors (miR-4459).

Thus, milk EV cargo is largely similar between donors while a few proteins and miRNAs are differentially expressed due to allergic sensitization. Although individual cargo has not met the significance thresholds after multiple testing correction, their cellular targets could be similar or act in concert, resulting in a shift in T cell modulation. To test this hypothesis, we performed *in silico* analysis and identified a total of 264 cellular targets in T cells, with 71 protein targets and 88 miRNA targets of non-allergic milk EVs, and 28 proteins and 76 miRNAs targets to allergic milk EVs. Using GO-analysis we determined the biological processes these targets were associated to, which included T cell activation, cytokine production, cell cycle and T cell signaling pathways/gene regulation (figure 2B). Interestingly, EVs from non-allergic milk regulate more T cell processes than EVs from allergic milk, except for PI3K-AKT signaling, suggesting that allergic milk EVs have a loss in T cell modulatory function.

### Target prediction of protein and miRNA cargo reveals an altered modulation of T cell proliferation and signaling

On basis of the functional data and the *in silico* modeling, we investigated how T cell proliferation was differentially targeted under influence of maternal sensitization. In our model the majority of miRNAs over-represented in allergic milk EVs target negative regulators of cell-cycle progression while those from non-allergic donors may target positive regulators of cell cycle, including cyclin-dependent kinases (Figure 3A). Similarly, we observed an over-representation in negative regulation of the JAK-STAT pathway by miRNAs from non-allergic mothers while for the PI3K-AKT pathway (that is downstream of CD28, growth factor receptor and integrin receptor) both positive and negative regulation by allergic and non-allergic EV cargo is possible (Figure 3A). In order to validate the *in silico* model, we cultured T cells in the presence of milk EVs and analyzed the expression of genes involved in T cell activation, anergy, and tolerance. In line with our proposed model, the transcription of 18 out of 28 selected genes (= 64%) was significantly less reduced during T cell activation in the presence of milk EVs derived from allergic compared to non-allergic mothers (supplementary Figure 4 and Figure 3B).

**Figure 3.**
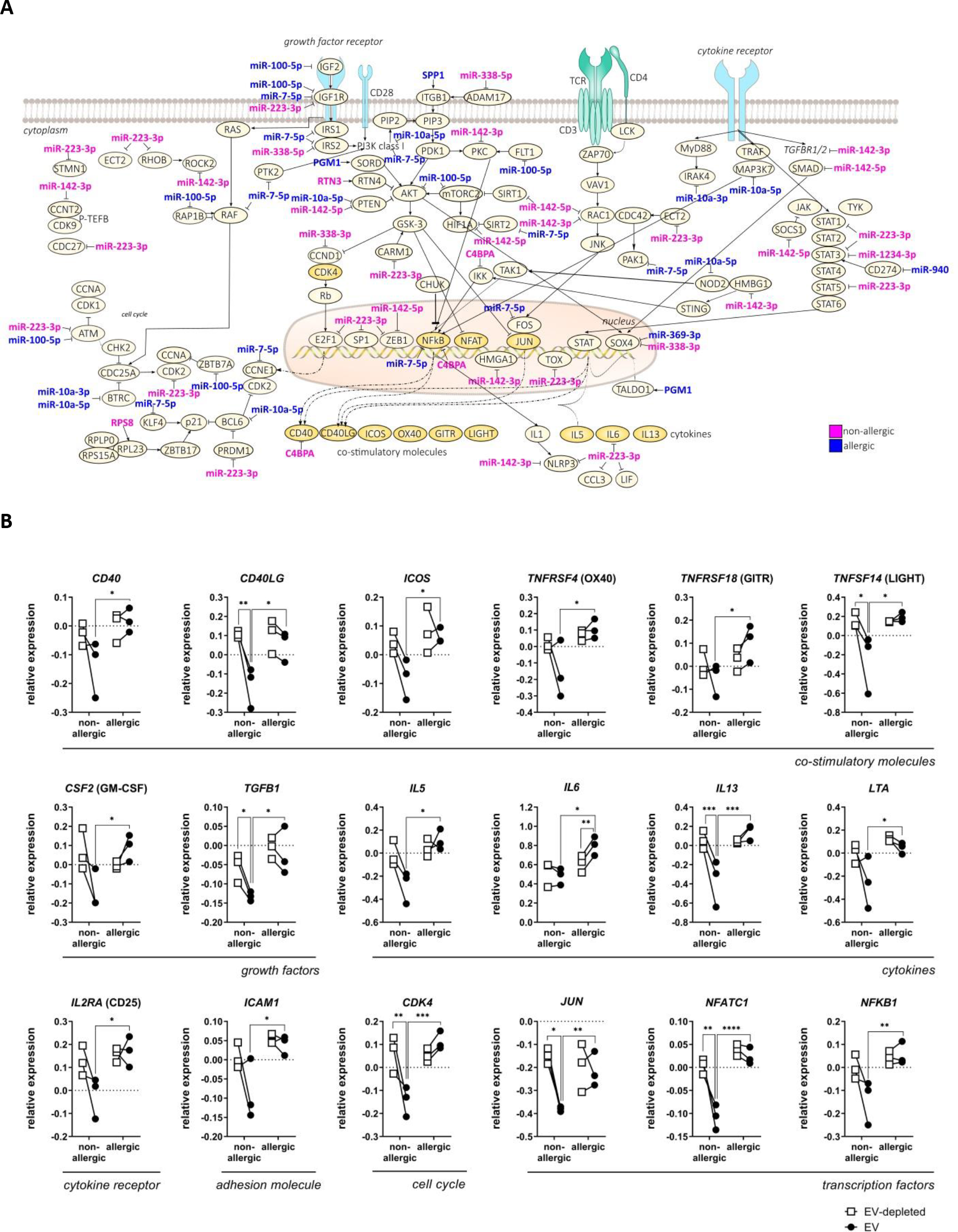
In silico analysis identifies cellular targets of milk EVs in cell cycle progression and T cell signaling pathways, and subsequent differentially modulated gene transcription. (A) Schematic representation of the cell cycle and T cell activation pathways like PI3K-AKT and JAK-STAT. The action of individual miRNAs and proteins (activating/interacting or inhibition) is depicted. (B) CD4+ T cells were stimulated with anti-CD3 and anti-CD28 in the presence of EVs or their EV-depleted controls from allergic (n=3) or non-allergic (n=3) mothers for 16 hours after which T cell RNA was extracted and gene expression analysis was performed. Gene expression was normalized to 4 housekeeping genes and delta Ct-values to anti-CD3 and anti-CD28 stimulated T cell controls were calculated and log-transformed (CD3/CD28 control values are 0). Shown are the paired results of each EV donor (●) with its EV-depleted control (□). Significance was calculated using repeated measures two-way ANOVA with Sidak’s multiple comparisons test and p-values defined as *p<0.05, **p<0.01, and **** p < 0.0001.

These genes included cyclin-dependent kinases (*CDK4)*, transcription factors (*JUN, NFKB1, NFATC1*), co-stimulatory (*CD40, CD40LG, ICOS, TNFRSF4, TNFRSF18, TNFSF14)* and adhesion molecules (*ICAM1*), growth factors (*CSF2, TGFB1*), cytokines (*IL5, IL6, IL13, LTA*), and cytokine receptors (IL2RA) (Figure 3B). Strikingly, IL-6 was not only differentially expressed between EVs, but also compared to the EV-depleted control (Figure 3B). Additionally, IL7R was induced in the presence of EV-depleted samples from allergic mothers. (Supplementary Figure 4), indicating the value of a procedural control in the accurate interpretation of EV-mediated effects [Théry]. Together, we show that that gene regulation by milk EVs is dependent on the allergic status of the mother and that different milk EV cargoes can act in concert to regulate T cell signaling pathways.

## Discussion

In this study we found that milk EVs of allergic donors were significantly less effective in suppressing T cell responses, suggesting that immune regulation in infants is altered due to maternal allergic sensitization.

No skewing of CD4+ T cells towards a Th1 or Th2 phenotype was seen under the influence of EVs. While increased transcription of *IL5, IL6*, and *IL13* could indicate a Th2 phenotype, only actual IL-6 secretion was augmented in the presence of milk EVs from allergic donors (Figure 2). This suggests that EVs from allergic mothers do not have the intrinsic capacity to skew T cell responses towards Th1 or Th2 immunity.

As EVs contain various bioactive cargoes that act in concert, the net-effect on the target cell will be determined by the collective functional EV cargo. Comparison of the protein and miRNA cargoes of EVs derived from allergic or non-allergic donors showed that the majority of proteins and miRNAs were similar in occurrence and abundance, which explains the overall inhibition of T cell activation induced by milk. Although no significant differences between groups were found after multi testing correction, synergistic effects of compositional alterations might actually drive the modest but significant effects on T cell inhibition. Furthermore, differences in post translational modifications or conformation of proteins, as well as differences in other bioactive EV-cargoes, e.g. mRNA, lncRNAs, metabolites or lipids might be fueling the differences in T cell inhibition.

To gain insights in the overall health and development of allergies in the infants from the lactating mothers participating in this study, we conducted a follow up study after 1 year of birth by questionnaire. Although we cannot exclude differences in allergic disorders appearing later in childhood, we did not find correlations to maternal allergies after 1 year, except for a disturbed sleep due to wheezing (supplementary Table 2).

Taken together, our study allows for a better understanding how maternal health affects T cell modulation and provides new insights into previously published reports whereby infants breastfed by sensitized mothers either had aggravated or attenuated allergy [Hamada, Leme, Mosconi, Baïz].

### Abbreviations

(EVs): Extracellular Vesicles
(ACCESS): Comparison of Human Milk Extracellular Vesicles in Allergic and Non-allergic Mothers
(EV-dpl): EV-depleted
(FASP): filter aided sample preparation
(LC-MS/MS): Liquid chromatography tandem mass spectrometry
(FDR): false discovery rate
(miRNA): microRNA
(PBMC): peripheral blood mononuclear cells
(CTV): CellTrace Violet
(CPM): counts per million

## Funding information and Acknowledgments

Subsidizing parties Nutricia Research and Technologiestichting STW as part of the partnership grant project 11676: Exosome-based biomarker profiling of breast milk: Definition of predictive immunemodulating biomarker profiles for the management of allergic disease development in infants, funded the research and the clinical Comparison of Human Milk Extracellular Vesicles in Allergic and Non-allergic Mothers (ACCESS) study. The research of MJCvH was further funded by the European Union’s Horizon 2020 Framework Programme under grant FETOPEN-801367 evFOUNDRY. AG, JJM, CO and MHMW were funded by The European Union’s Horizon 2020 research and innovation programme under the Marie Skłodowska-Curie grant agreement No 722148. TAPD was supported by the European Research Council under the European Union’s Seventh Framework Programme [FP/2007-2013]/ERC Grant Agreement number [337581].

We thank Hailiang Mei, Wai-Yi Leung (Sequencing Analysis Support Core, Leiden University Medical Center, Leiden, The Netherlands) and Henk P.J. Buermans (Department of Human Genetics and Leiden Genome Technology Center, Leiden University Medical Center, Leiden, The Netherlands) for their contributions to miRNA analysis.

## Supplementary methods

### High Resolution Flow Cytometry

PKH67 labeling and high-resolution flow cytometric analysis was performed as previously described [Kleinjan 2021]: Pelleted samples of the pooled density fractions were resuspended in 40 μl PBS, of which 10 μl of resuspended EV samples were fluorescently labeled with PKH67 (Sigma-Aldrich) and separated from unbound PKH67 and protein aggregates by overnight density gradient centrifugation at 192,000 × g for 15–18 h at 4◦C using an SW40 rotor, according to the previously described protocols (34, 35). From the top of the tube, 12 fractions of 1 mL were collected in Eppendorf tubes (Eppendorf) and stored at 4◦C until further analysis. High-resolution flow cytometric analysis of PKH67-stained samples was performed by fluorescent threshold triggering on a BD Influx cell sorter (BD Biosciences) that was dedicated to and optimized for detection of submicron-sized particles. Detailed descriptions of both the hardware adaptations and methods used have been previously published (34, 36, 37). To exclude detection of EV swarms, serial dilutions of samples were performed. All 12 fractions from the second density gradient were then individually measured at optimal dilution. Data analysis was performed with FlowJo version 10.5.0. (FlowJo) To determine the total number of PKH67-labeled particles present in the pooled density gradient fractions 4–6 and 7–9 derived from the first gradient, the sum total of PKH67-positive events in fractions 3–12 of the second density gradient was calculated and corrected with the dilution factor. Fractions 1–2 were omitted, because these contained unbound PKH67 dye aggregates, not EVs.

### CD4+ T cell isolation and stimulation

T cells were isolated and stimulated as described in the methods. Cells were cultured for 6 days in medium alone, or with a dilution series of EVs, or EV-depleted control (750 μl, 350 μl, 188 μl, 94 μl) supplemented with medium to come to a total volume of 1 ml. Proliferation was assessed by labeling cells with 2 μM CellTrace Violet (Invitrogen) prior to culture. Cells were acquired on a BD Canto II (BD Bioscience), and analyzed by FlowJo (V10.1). From comparison in T cell modulatory function, from each group one donor was selected according to total number of counts from the high-resolution flow cytometry analysis, where the non-allergic donor had 449,458 events and the allergic donor 164,117 events (2,74x difference).

## Supplementary Figures

**Supplementary Figure 1:**
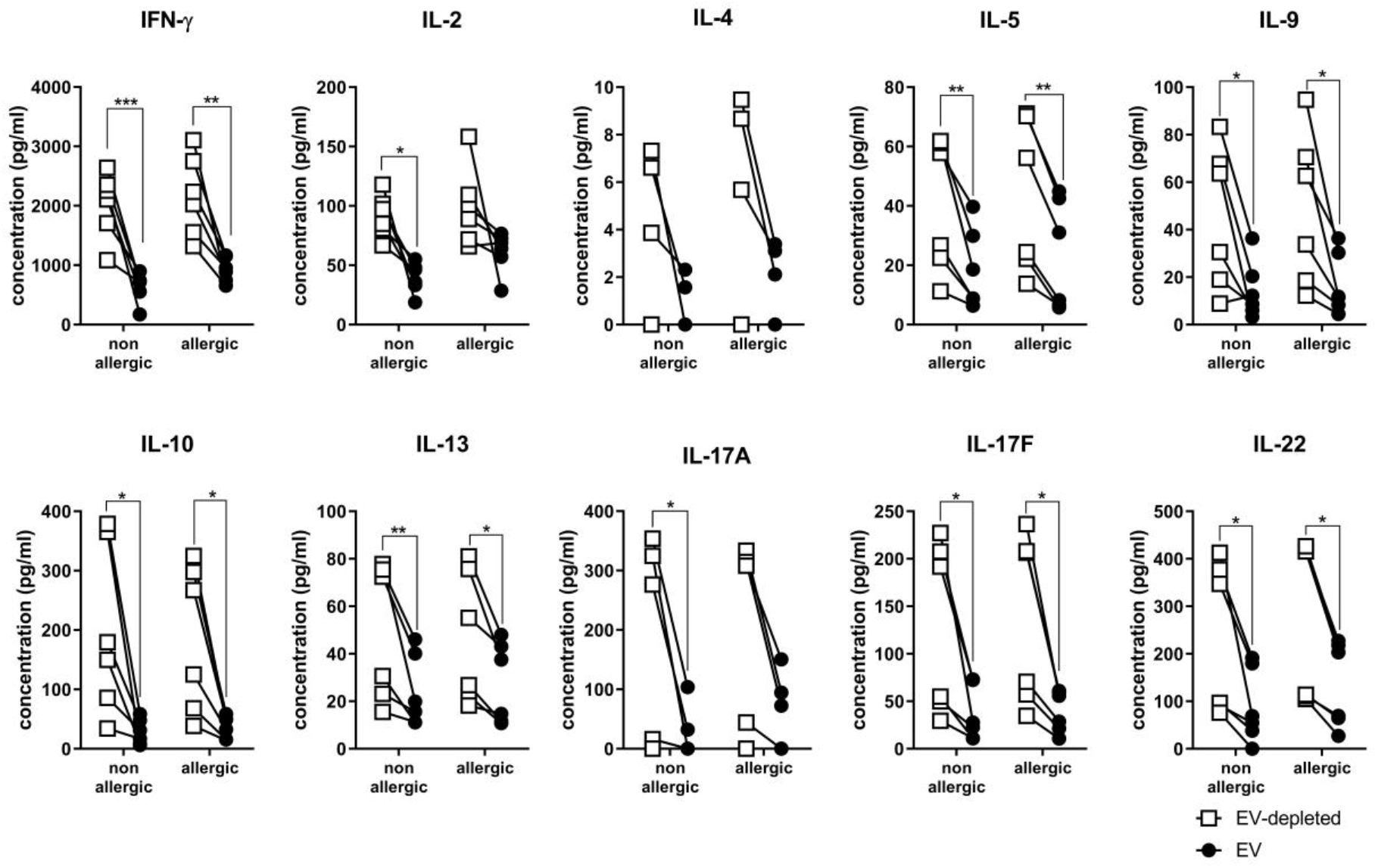
Human milk EVs from non-allergic and allergic mothers inhibit secretion of various Th1/2/9/17 related cytokines by CD4+ T cells. CD4+ T cells of 2 different donors were stimulated with anti-CD3 and anti-CD28 in the presence EVs (●) or its EV-depleted control (□) from allergic (n=3) or non-allergic (n=3) mothers. Supernatants were harvested and analyzed for cytokines after 2 and 6 days of culture. Except for IL-2 and IL-4, data of day 6 is shown. Choice for day 2 or day 6 was made depending on when the optimal resolution between EV of allergic or non-allergic donors was obtained. Graphs show the paired results obtained for each EV donor and its EV-depleted control in pg/ml. Significance was calculated using repeated measures two-way ANOVA with Sidak’s multiple comparisons test. P-values are defined as * < 0.05, **p < 0.01 and ***p < 0.001.

**Supplementary Figure 2:**
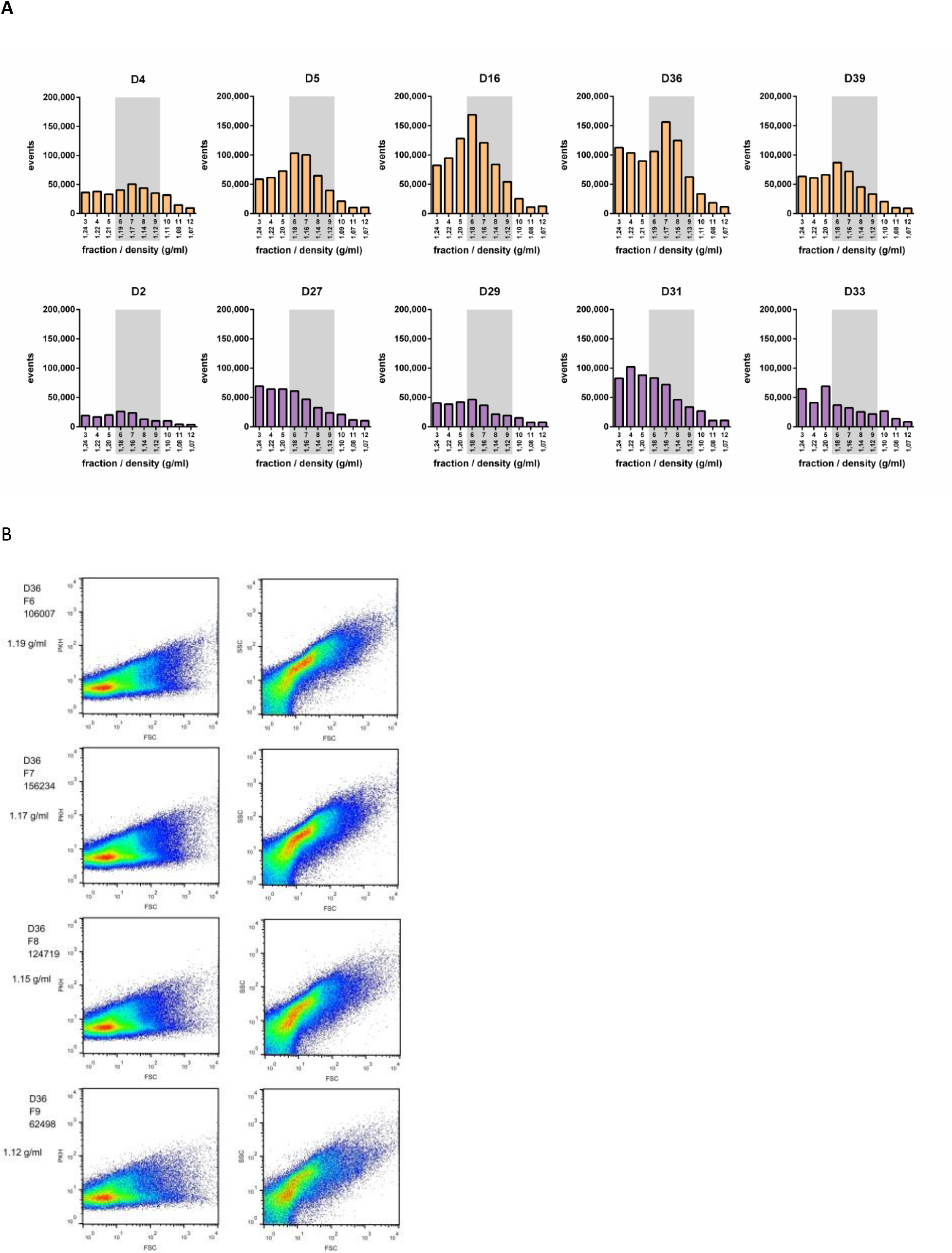
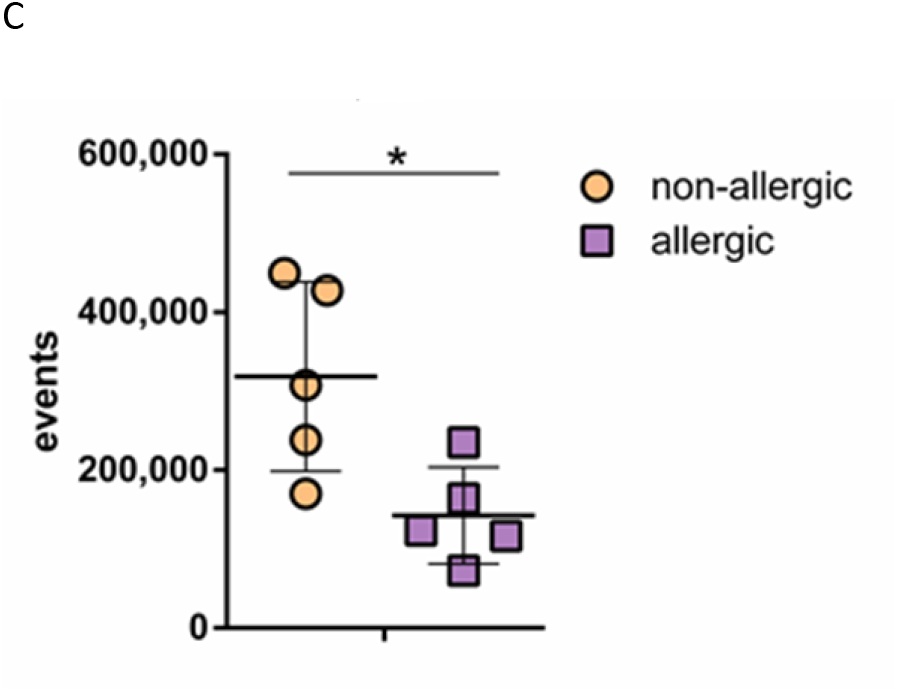
High Resolution Flow Cytometry. Quantitative EV analysis of raw and differently processed bovine milk using single-particle EV-based high-resolution flow cytometry. A) Quantitative high-resolution flow cytometric analysis of EVs showing the total number of PKH67-labeled events per fraction with the EV-enriched fractions highlighted in grey (fractions 6-9, 1.19-1.12 g/ml). Fractions 1–2 were omitted, because these contained unbound PKH67 dye aggregates, not EVs. B) Typical scatterplots of two representative donors from each group showing PKH67 fluorescence and FSC, as well as SSC and FSC, of PKH67-positive EVs in EV-enriched density fractions. C)To determine the total number particles, the sum total of PKH67-positive events in fractions 6-9 was calculated and corrected with the dilution factor. Significance was calculated using paired t test and p-values defined as *p<0.05.

**Supplementary Figure 3:**
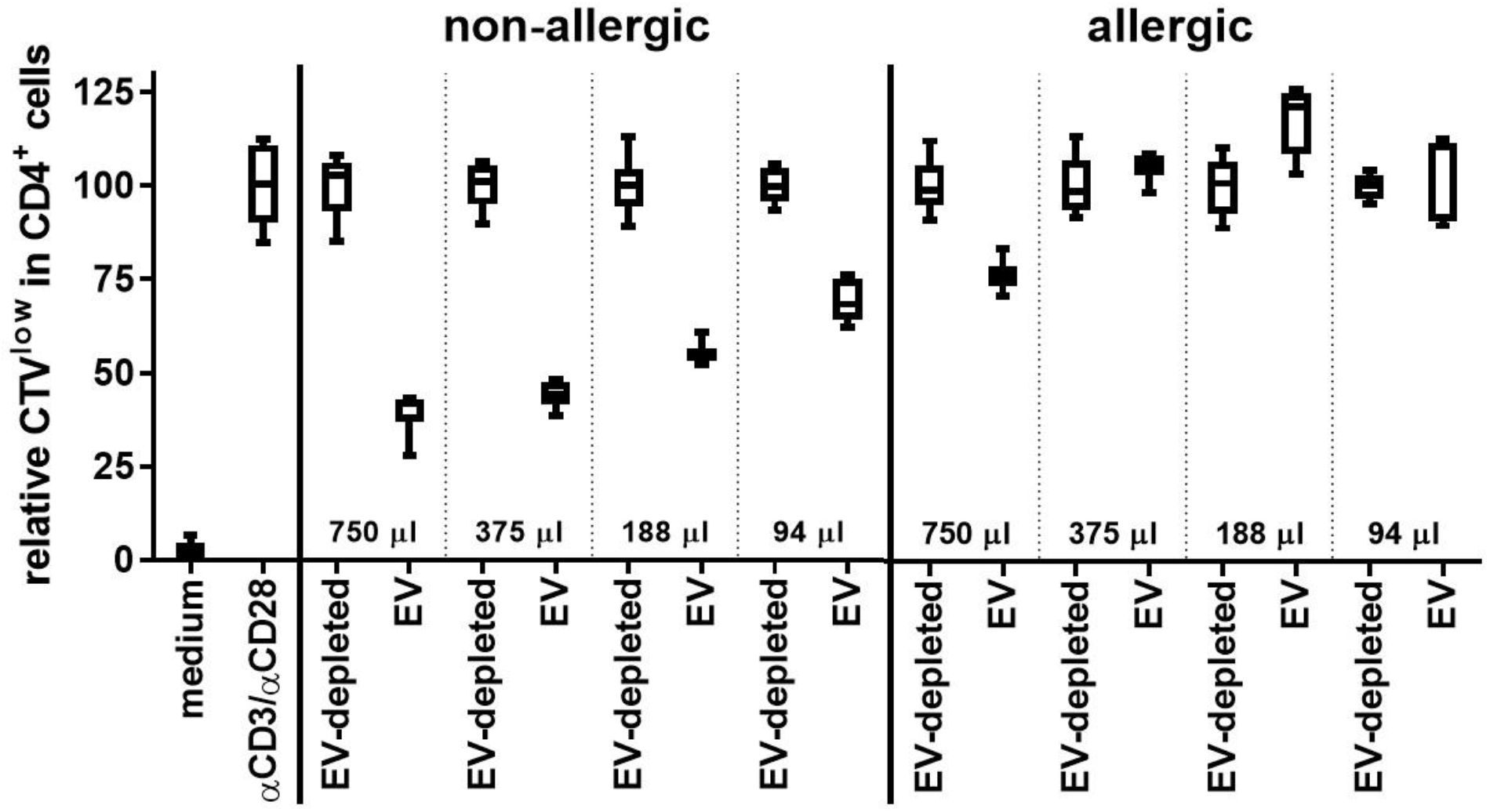
Inhibition of CD4+ T cell activation is reduced by milk EVs from an allergic mother even at high concentrations, while low volumes of milk EVs from a non-allergic mother already suppress T cell activation. CD4+ T cells were stimulated with anti-CD3 and anti-CD28 in the presence of EVs or its EV-depleted control sample from an allergic (n=1) or non-allergic (n=1) mother. For flow cytometry, T cells from 1 donors were labeled with CellTrace Violet (CTV), cultured for 6 days, harvested and analyzed. Quantification of proliferation is shown for cells which had undergone one or more proliferation cycles (and were thus lower in CTV signal intensity than non-proliferating T cells). Graphs show boxplots for each culture condition (triplicate samples) with relative expression of CTVlow CD4+ T cells following incubation with the indicated conditions, with the average of each paired EV-depleted control set to 100%. According to high-resolution flow cytometry, the allergic milk donor had 2,74 times less EVs in her milk than the non-allergic milk donor, thus 274 µl EV sample from the non-allergic donor (which is in between the 375µl and 188µl condition) would be similar in EV counts as 750 µl EV sample of the allergic donor.

**Supplementary Figure 4:**
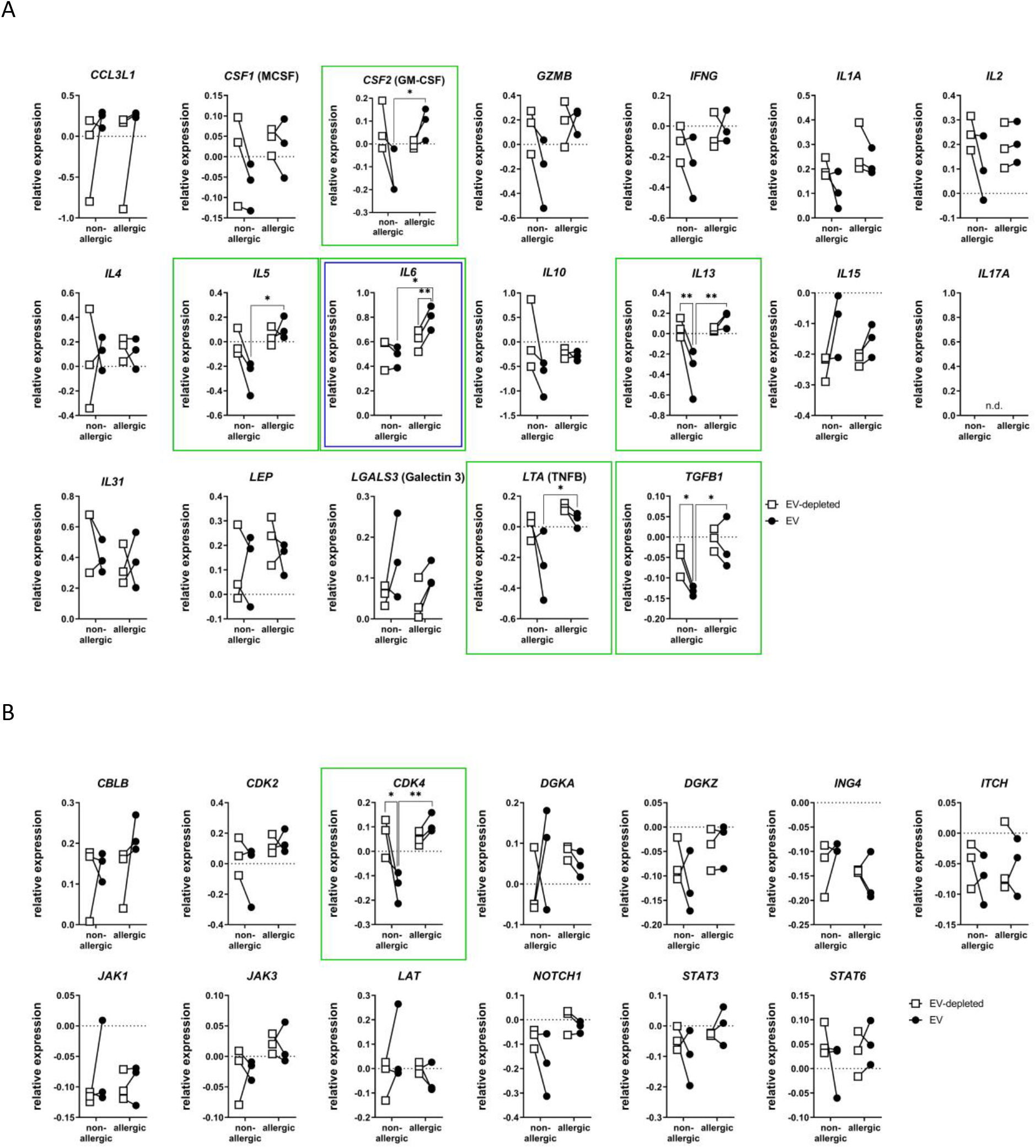
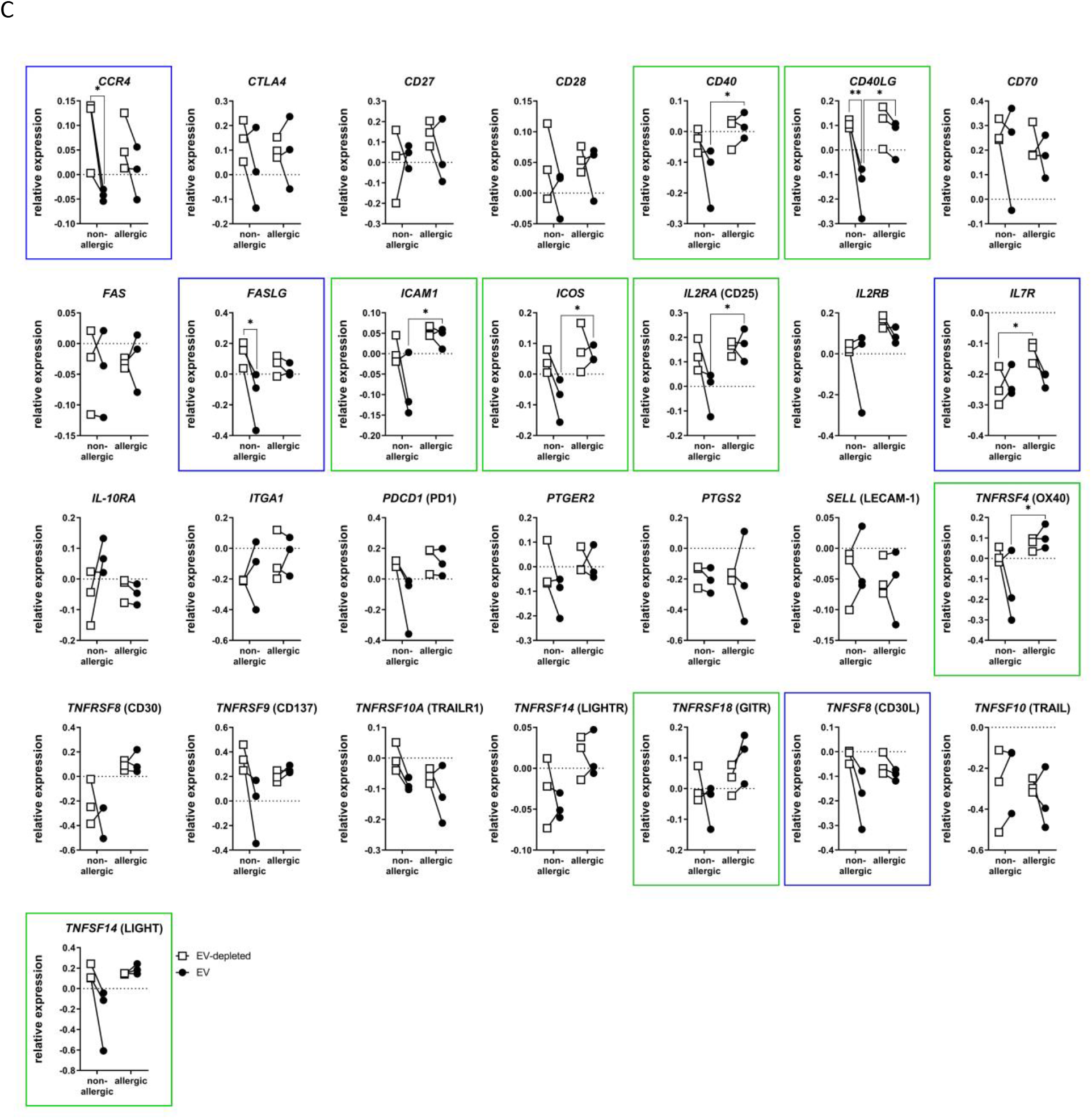
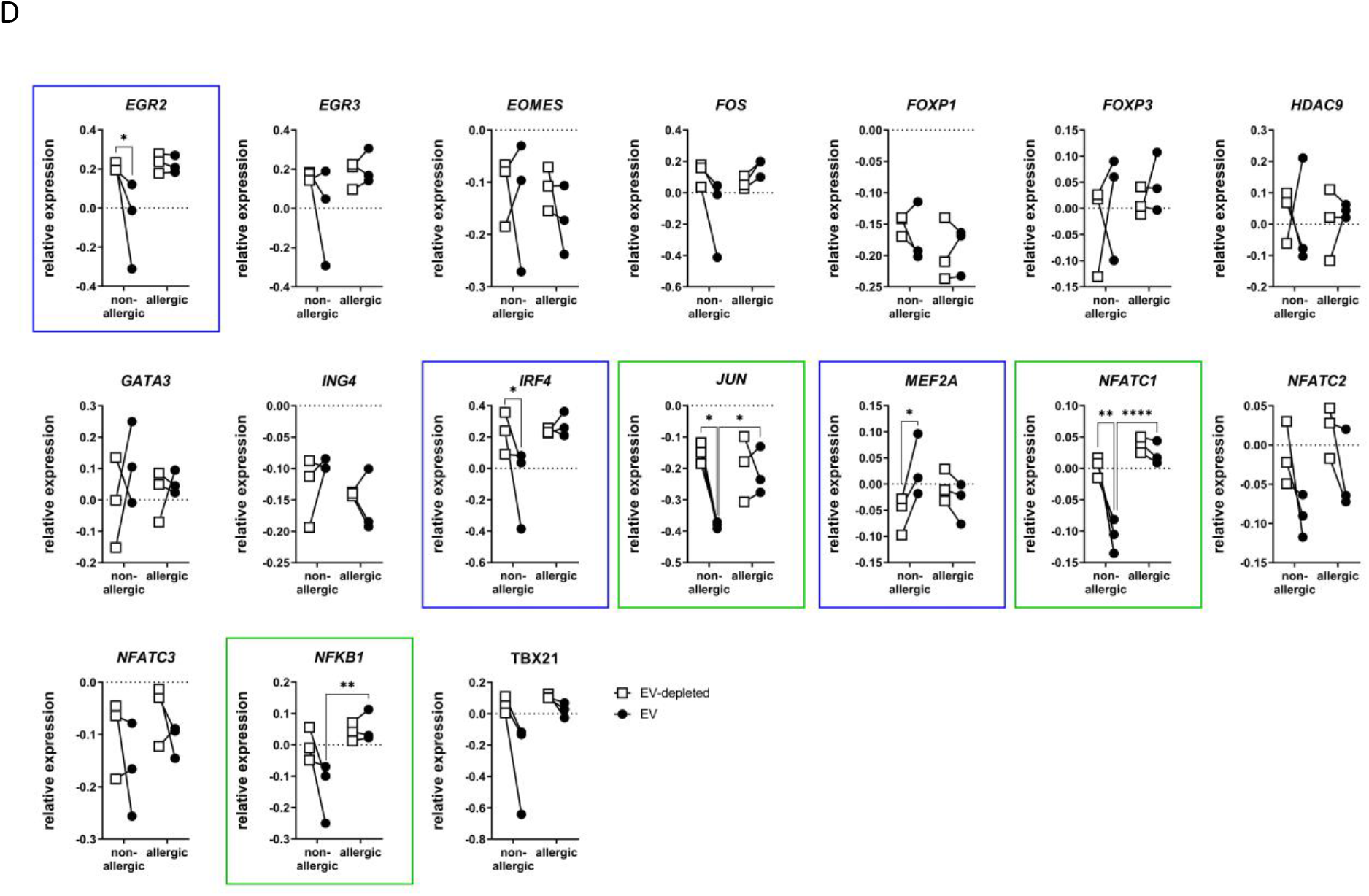
Gene expression analysis of CD4+ T cells stimulated with CD3/CD28 in the presence of EV or EV-depleted controls from allergic or non-allergic mothers. CD4+ T cells were stimulated with anti-CD3 and anti-CD28 in the presence of EVs (●) or their EV-depleted controls (□) from allergic (n=3) or non-allergic (n=3) mothers for 16 hours after which T cell RNA was extracted and gene expression analysis was performed. Gene expression was normalized to 4 housekeeping genes and delta Ct-values to CD3/CD28 stimulated T cell controls were calculated and log-transformed (CD3/CD28 control values are 0). Shown are the paired results of each EV donor with its EV-depleted control. Shown is the gene expression analysis of (A) secretory mediators, (B) Intracellular signaling molecules, (C) extracellular receptors and ligands, and (D) transcription factors. Green boxes indicate genes which were found to significantly differ between EV from allergic and non-allergic mothers, as presented in Figure 3. Blue boxes indicate that significant differences were detected in conditions other than allergic and non-allergic EV, like EV vs. EV-depleted. N.d. Not detected (IL17A). Significance was calculated using repeated measures two-way ANOVA with Sidak’s multiple comparisons test and p-values defined as *p<0.05, **p<0.01, and **** p < 0.0001.

## Supplementary Tables

**Supplementary Table 1.**
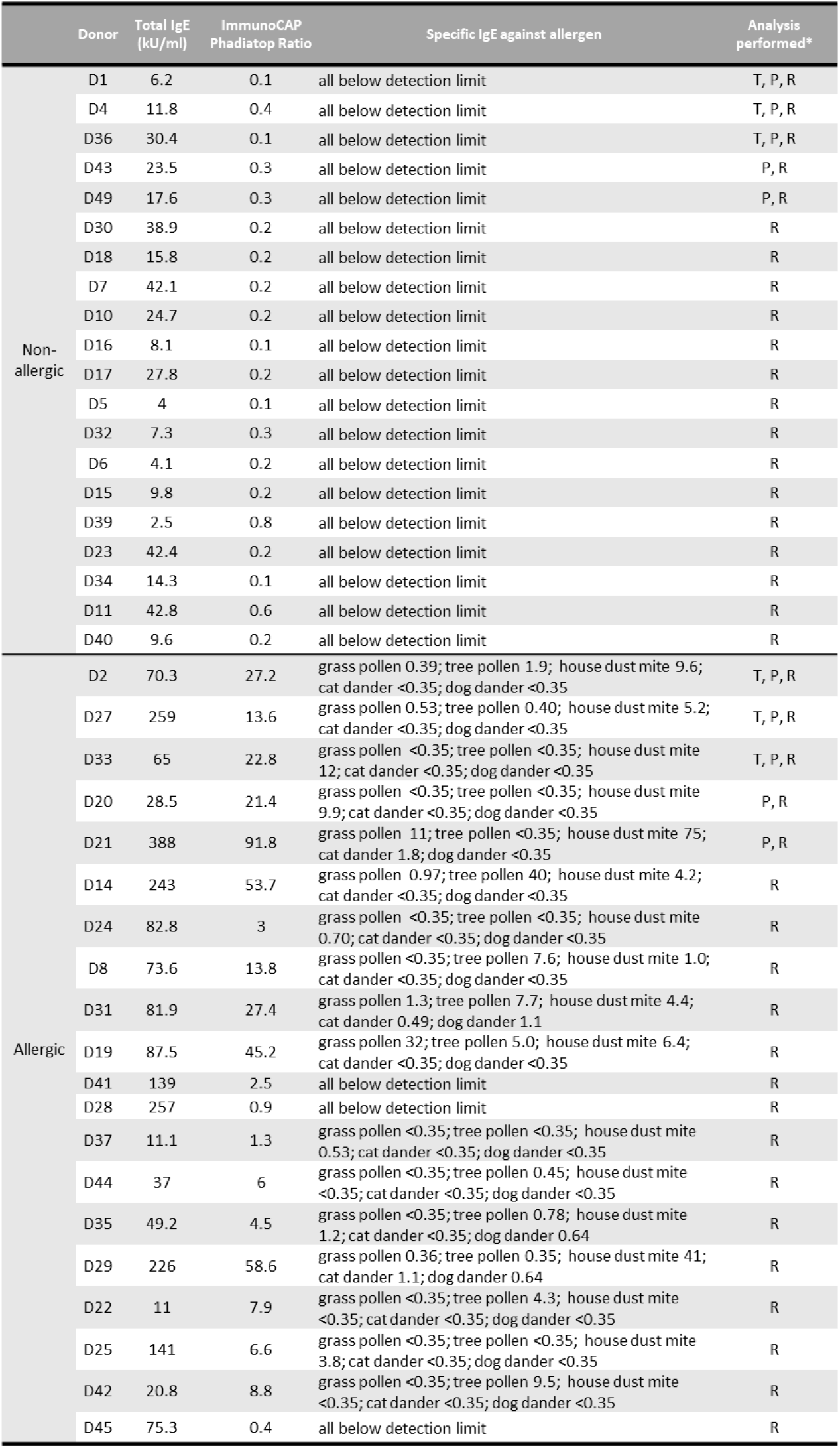
Data of individual donors used in the present study. Table lists donor ID according the ACCESS study register, whether donors were classified as non-allergic or allergic, total IgE (kU/L), ImmunoCap Phadiatop ratios, and specific IgE. The last column lists whether donors were included in functional T cell assays (T), proteomics analysis (P) and/or RNA deep sequencing (R).

|  | Donor | Total IgE<br>(kU/ml) | ImmunoCAP<br>Phadiatop Ratio | Specific IgE against allergen | Analysis<br>performed* |
| --- | --- | --- | --- | --- | --- |
| Non-allergic | D1 | 6.2 | 0.1 | all below detection limit | T, P, R |
|  | D4 | 11.8 | 0.4 | all below detection limit | T, P, R |
|  | D36 | 30.4 | 0.1 | all below detection limit | T, P, R |
|  | D43 | 23.5 | 0.3 | all below detection limit | P, R |
|  | D49 | 17.6 | 0.3 | all below detection limit | P, R |
|  | D30 | 38.9 | 0.2 | all below detection limit | R |
|  | D18 | 15.8 | 0.2 | all below detection limit | R |
|  | D7 | 42.1 | 0.2 | all below detection limit | R |
|  | D10 | 24.7 | 0.2 | all below detection limit | R |
|  | D16 | 8.1 | 0.1 | all below detection limit | R |
|  | D17 | 27.8 | 0.2 | all below detection limit | R |
|  | D5 | 4 | 0.1 | all below detection limit | R |
|  | D32 | 7.3 | 0.3 | all below detection limit | R |
|  | D6 | 4.1 | 0.2 | all below detection limit | R |
|  | D15 | 9.8 | 0.2 | all below detection limit | R |
|  | D39 | 2.5 | 0.8 | all below detection limit | R |
|  | D23 | 42.4 | 0.2 | all below detection limit | R |
|  | D34 | 14.3 | 0.1 | all below detection limit | R |
|  | D11 | 42.8 | 0.6 | all below detection limit | R |
|  | D40 | 9.6 | 0.2 | all below detection limit | R |
| Allergic | D2 | 70.3 | 27.2 | grass pollen 0.39; tree pollen 1.9; house dust mite 9.6;<br>cat dander <0.35; dog dander <0.35 | T, P, R |
|  | D27 | 259 | 13.6 | grass pollen 0.53; tree pollen 0.40; house dust mite 5.2;<br>cat dander <0.35; dog dander <0.35 | T, P, R |
|  | D33 | 65 | 22.8 | grass pollen <0.35; tree pollen <0.35; house dust mite<br>12; cat dander <0.35; dog dander <0.35 | T, P, R |
|  | D20 | 28.5 | 21.4 | grass pollen <0.35; tree pollen <0.35; house dust mite<br>9.9; cat dander <0.35; dog dander <0.35 | P, R |
|  | D21 | 388 | 91.8 | grass pollen 11; tree pollen <0.35; house dust mite 75;<br>cat dander 1.8; dog dander <0.35 | P, R |
|  | D14 | 243 | 53.7 | grass pollen 0.97; tree pollen 40; house dust mite 4.2;<br>cat dander <0.35; dog dander <0.35 | R |
|  | D24 | 82.8 | 3 | grass pollen <0.35; tree pollen <0.35; house dust mite<br>0.70; cat dander <0.35; dog dander <0.35 | R |
|  | D8 | 73.6 | 13.8 | grass pollen <0.35; tree pollen 7.6; house dust mite 1.0;<br>cat dander <0.35; dog dander <0.35 | R |
|  | D31 | 81.9 | 27.4 | grass pollen 1.3; tree pollen 7.7; house dust mite 4.4;<br>cat dander 0.49; dog dander 1.1 | R |
|  | D19 | 87.5 | 45.2 | grass pollen 32; tree pollen 5.0; house dust mite 6.4;<br>cat dander <0.35; dog dander <0.35 | R |
|  | D41 | 139 | 2.5 | all below detection limit | R |
|  | D28 | 257 | 0.9 | all below detection limit | R |
|  | D37 | 11.1 | 1.3 | grass pollen <0.35; tree pollen <0.35; house dust mite<br>0.53; cat dander <0.35; dog dander <0.35 | R |
|  | D44 | 37 | 6 | grass pollen <0.35; tree pollen 0.45; house dust mite<br><0.35; cat dander <0.35; dog dander <0.35 | R |
|  | D35 | 49.2 | 4.5 | grass pollen <0.35; tree pollen 0.78; house dust mite<br>1.2; cat dander <0.35; dog dander 0.64 | R |
|  | D29 | 226 | 58.6 | grass pollen 0.36; tree pollen 0.35; house dust mite 41;<br>cat dander 1.1; dog dander 0.64 | R |
|  | D22 | 11 | 7.9 | grass pollen <0.35; tree pollen 4.3; house dust mite<br><0.35; cat dander <0.35; dog dander <0.35 | R |
|  | D25 | 141 | 6.6 | grass pollen <0.35; tree pollen <0.35; house dust mite<br>3.8; cat dander <0.35; dog dander <0.35 | R |
|  | D42 | 20.8 | 8.8 | grass pollen <0.35; tree pollen 9.5; house dust mite<br><0.35; cat dander <0.35; dog dander <0.35 | R |
|  | D45 | 75.3 | 0.4 | all below detection limit | R |

**Supplementary Table 2.** Table lists various information on the infants health during or after 1 year of life. Significance was calculated using Fisher exact test and only ‘disturbed sleep due to wheezing’ occurred significantly more in infants breastfed by allergic mothers. Significance was calculated using Fisher exact test.

| Comparison of groups | non-allergic<br>(n=29) | allergic<br>(n=30) | P value |
| --- | --- | --- | --- |
| maternal perception infants health: very healthy | 8/29 | 8/30 | 0.9999 |
| maternal perception infants health: sometimes ill | 21/29 | 17/30 | 0.1881 |
| maternal perception infants health: regularly ill | 0/29 | 5/30 | 0.0522 |
| infant ever had a variably present itching rash | 10/29 | 8/30 | 0.5796 |
| this rash dissapeared completely | 8/10 | 5/7 | 0.9999 |
| infant woken up due to rash | 3/10 | 0/7 | 0.2279 |
| infant suffered from wheezing | 7/29 | 8/30 | 0.9999 |
| 1 to 3 episodes of wheezing | 6/7 | 5/8 | 0.5692 |
| 4 to 12 episodes of wheezing | 1/7 | 3/8 | 0.5692 |
| **disturbed sleep due to wheezing** | **2/7** | **7/8** | **0.0406** |
| infant suffered from a dry cough without having a cold | 2/29 | 7/30 | 0.1455 |
| suffered from a cough during the day | 22/29 | 23/30 | 0.9999 |
| once or more coughs during the day | 12/22 | 14/23 | 0.7666 |
| suffered from low-pitched wheezes or coarse crackles | 7/29 | 8/30 | 0.9999 |
| suffered from a hortness of breath | 4/29 | 9/30 | 0.2092 |
| once or more shortness of breath | 2/4 | 5/9 | 0.9999 |
| infant diagnosed with asthma | 0/29 | 1/30 | 0.9999 |
| suffered from running or stuffy nose | 3/29 | 8/30 | 0.1806 |
| ever had itching eyes | 6/29 | 7/30 | 0.2092 |

